# Image-based quantification of *Arabidopsis thaliana* stomatal aperture from leaf images

**DOI:** 10.1101/2022.11.30.518467

**Authors:** Momoko Takagi, Rikako Hirata, Yusuke Aihara, Yuki Hayashi, Miya Mizutani-Aihara, Eigo Ando, Megumi Yoshimura-Kono, Masakazu Tomiyama, Toshinori Kinoshita, Akira Mine, Yosuke Toda

## Abstract

The quantification of stomatal pore size has long been a fundamental approach to understand the physiological response of plants in the context of environmental adaptation. Automation of such methodologies not only alleviates human labor and bias, but also realizes new experimental research methods through massive analysis. Here, we present an image analysis pipeline that automatically quantifies stomatal aperture of *Arabidopsis thaliana* leaves from brightfield microscopy images containing mesophyll tissue as noisy backgrounds. By combining a YOLOX-based stomatal detection submodule and a U-Net-based pore segmentation submodule, we achieved 0.875 mAP_50_ (mean average precision; stomata detection performance) and 0.745 IoU (intersection of union; pore segmentation performance) against images of leaf discs taken with a brightfield microscope. Moreover, we designed a portable imaging device that allows easy acquisition of stomatal images from detached/undetached intact leaves on-site. We further combined this device with fine-tuned models of the pipeline we generated here and recapitulated manual measurement of stomatal responses against pathogen inoculation. Utilization of our hardware and pipeline for automated stomatal aperture measurements is expected to accelerate research on stomatal biology of model dicots.

## Introduction

Stomata, which consist of a pair of guard cells, control gas exchange between the leaf and the atmosphere. In response to various environmental stimuli, plants regulate stomatal aperture for adaptation. Plants perceive blue light as a signal that triggers K^+^ ion uptake in guard cells; plants also use red light as an energy source for photosynthesis in chloroplasts, resulting in stomatal opening for CO_2_ uptake (Inoue and Kinoshita, 2017; Matthews et al., 2020; Shimazaki et al., 2007). Plants synthesize the phytohormone abscisic acid (ABA) under drought conditions to promote stomatal closure and thereby prevent excess water loss (Hsu et al., 2021). In addition to abiotic stimuli, stomata also respond to biotic stimuli. For example, plants close their stomata by detecting cues from bacterial and fungal infections. To counter this immune response, the phytopathogenic bacterium *Pseudomonas syringae* pv. *tomato (Pst*) DC3000 and the fungus *Fusicoccum amygdali* produce coronatine and fusicoccin, respectively, to manipulate the plant stomatal signaling pathway to force stomata open to ensure a physical entry point for infection (Camoni et al., 2019; Melotto et al., 2017).

To elucidate the mechanisms underlying the physiology of guard cell regulation, quantification of stomatal aperture from microscopy observations has been commonly used as a metric. Traditional observation methods use an eyepiece micrometer to measure stomatal aperture from either epidermal peels, epidermal fractions prepared using a blender, or partial leaf fragments, either prepared by forceps or hole punches (leaf discs). However, the number of experimental conditions that can be performed at once has been restricted because the manual quantification of numerous stomatal pores is labor-intensive. Instead, alternative metrics have been measured as proxy to estimate the degree of stomatal aperture, including thermal imaging as an estimation of leaf transpiration (Hashimoto et al., 1984), measuring transpiration rate with a porometer (e.g. Delta-T AP4 Porometer; Delta-T Devices, UK), or utilizing fluids with varying viscosity (Hack, 1974). Importantly, direct quantification of stomatal aperture is still of interest to researchers, as it provides rich and direct information related to the sensitive dynamics of stomatal movements. In such cases, ImageJ and Fiji (Schindelin et al., 2012; Schneider et al., 2012) are some of the most commonly used software for biologists when quantifying stomatal aperture from individually collected images in specific conditions. Recently, owing to technical advances in image analysis libraries, image-based automatic quantification has gradually been introduced to the community over the past decade.

Several groups have implemented image analysis pipelines using confocal microscopy images. For example, stomatal apertures of Arabidopsis (*Arabidopsis thaliana*) have been quantified from such microscopy-assisted capture using a fluorescent actin marker (Shimono et al., 2016), cell wall autofluorescence (Bourdais et al., 2019), or a fluorescent dye (Eisele et al., 2016; Higaki et al., 2014), each differently highlighting stomatal pore areas within the images. While these approaches can greatly reduce background noise and facilitate the application of efficient analysis modules, the cost of both maintaining and running a confocal microscopy facility, as well as sample preparation, can be a potential obstacle against routine application. Notably, using a benchtop brightfield microscope or an equivalent observation device is expected to alleviate the above-mentioned issues. However, high noise in the acquired images (e.g., uneven light irradiation due to sample thickness, appearance of non-stomatal components such as pavement cell and mesophyll cells, and partially out-of-focus images) in turn limits the deployment of such devices.

Recently, incorporating machine learning into the image analysis pipeline has aided the development of automatic quantification of stomatal aperture. For example, stomatal detection based on histogram-of-oriented-gradients (HOG) followed by image skeletonization and ellipse fitting was applied to grapevine (*Vitis vinifera*) images (Jayakody et al., 2017). Similarly, HOG and convolutional neural network (CNN) were combined to detect and discriminate between open and closed stomata, along with morphological shape filtering for pore quantification in Benghal dayflower (*Commelina benghalensis*) (Toda et al., 2018). Stomatal detection by Faster Region-based CNN (Faster R-CNN) and ChanVese-based pore segmentation was applied for poplars (*Populus* L.) (Li et al., 2019). Mask R-CNN was implemented to both count stomata and quantify the stomatal aperture of black poplar (*Populus nigra*) and gingko (*Gingko biloba*) (Song et al., 2020) or epidermal peels of Arabidopsis and barley (*Hordeum vulgare*) (Sai et al., 2022).

As real-time observations of stomata from intact leaves allow for a rigorous evaluation of the physiological response of plants at specific moments, methods that allow such observations have been in high demand by many researchers. Clark 2019 created a custom chamber to fix the leaves of *Tradescantia spathacea* to the stage of an upright microscope for analysis of stomatal dynamics in response to light and CO_2_ levels. Similarly, the VHX-2000 (Keyence, Japan), a microscope-based apparatus with a large depth of field, was used to acquire a fully focused image of black poplar leaves and quantify stomatal aperture (Song et al., 2020). Several research groups have attempted to utilize or produce a portable imaging device. For example, the handheld microscope ProScope HR2 (Bodelin technologies, USA) was used to quantify stomatal aperture of maize (*Zea mays*) leaves in an undetached intact state (Liang et al., 2022). Finite 40× objective lenses were used to assemble a customized portable microscope, allowing the monitoring of stomatal movements from tomato (*Solanum lycopersicum*) plants grown in the field (Purwar and Lee, 2019). The resulting acquired images were processed with a customized image analysis pipeline or by manual quantification to evaluate stomatal aperture.

In this study, we aimed to develop an image analysis module that can quantify stomatal aperture of Arabidopsis leaves to accelerate routine benchtop analysis involved in physiological research. Up-to-date methods to quantify Arabidopsis stomatal aperture from brightfield microscope images have been limited, with the exception of a previous study (Sai et al., 2022). Notably, the module built in the Sai et al. study was specifically designed for epidermal peels. Since epidermal peels consist of a single cell layer (guard cells and pavement cells) with almost noiseless background, the stomatal pores in the acquired images are clear, allowing for the precise quantification of stomatal dynamics. In addition to guard cell autonomous signals, recent findings indicate that signals derived from mesophyll cells also have an important role in the regulation of stomatal movements (Matthews et al., 2020). Thus, quantification of stomatal aperture from materials with mesophyll cells still attached (i.e., undetached intact leaves, detached leaves, or leaf discs) has been a subject of interest. However, the difficulty of image focusing due to the bumpy leaf surface as well as the greater noise emanating from the mesophyll tissue has been a technical challenge for the development of an efficient image analysis methodology.

Here, we established an image analysis pipeline to process such noisy leaf images. By combining a YOLOX-based stomata detection and U-Net-based stomatal pore segmentation submodules, we achieved a mean average precision value (mAP_50_) of 0.875 and an intersection of union (IoU) of 0.745, which accounts to a stomatal aperture (pore width) quantification error of 0.2 ± 0.2 μm compared to manual measurements when evaluated against detached leaf disc images taken with an upright brightfield microscope. Moreover, we designed a portable device that allows easy stomatal observations from both undetached intact and detached leaves. We further fine-tuned the models of our image analysis pipeline for their use with images captured with our new portable device and successfully recapitulated manual measurement of stomatal responses against *Pst* DC3000 infection. Collectively, we present a system that can automatically evaluate the response of stomatal regulation of Arabidopsis leaves in various situations.

## Results

### Dataset and Machine Learning Model Construction

We constructed a two-step image analysis pipeline to quantify stomatal aperture (Fig. 1A). The first object detection module identifies and retrieves stomatal coordinates, simultaneously classifying each stoma as opened or closed. The second semantic segmentation module processes subimages from all open stomata and extracts stomatal pore area. The profiles of measured stomatal pores (e.g., area, width, width/length ratio) are aggregated with the results from closed stomata (with apertures of 0 μm) before being used for further analysis. This pipeline allows the extraction of the stomatal aperture averages or medians along with the visualized output (Fig. 1B).

**Figure 1.**
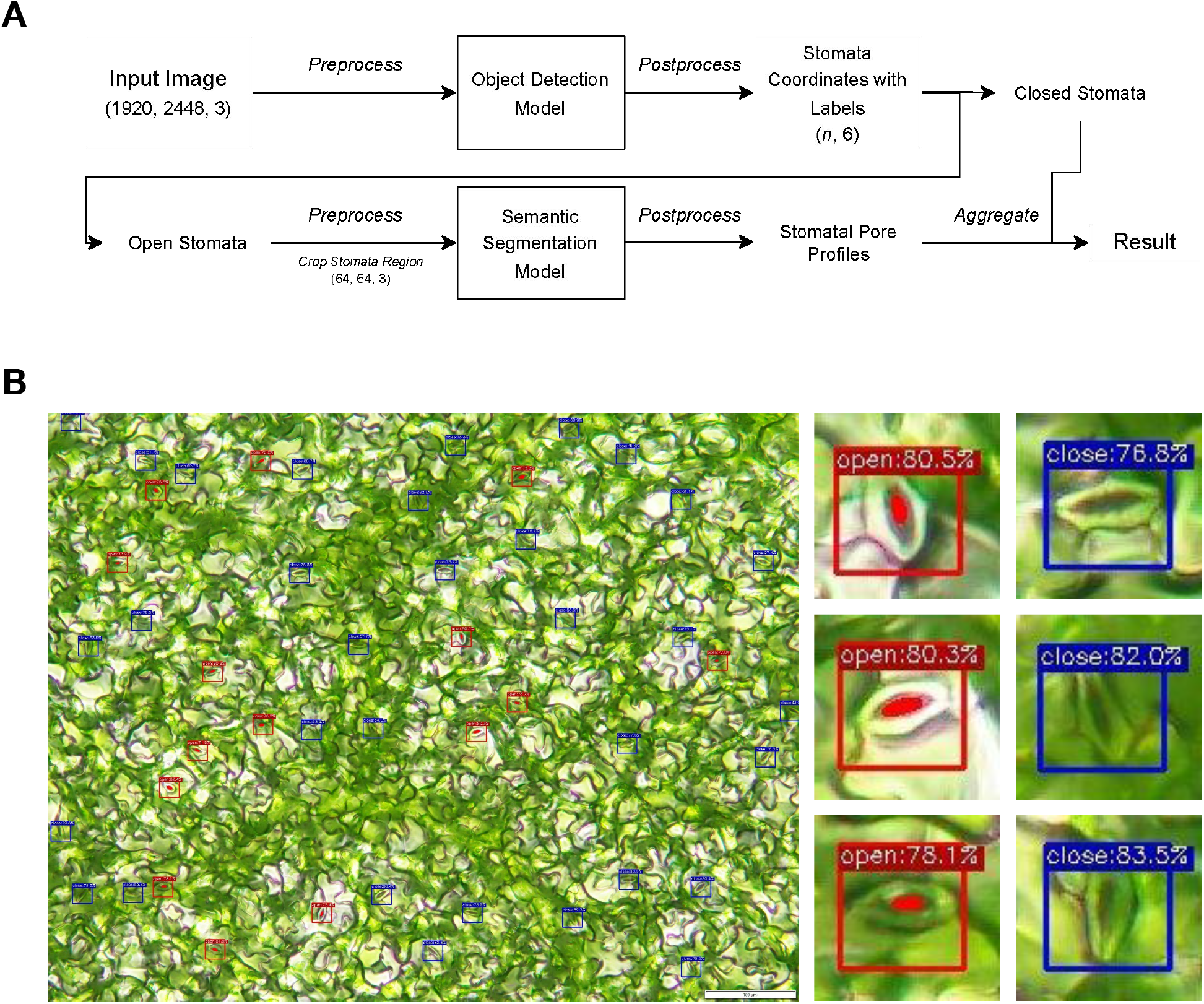
Schematic diagram of the stomatal aperture quantification pipeline and representative output. (A) Schematic diagram of the quantification pipeline for Arabidopsis stomatal pores. Numbers in parentheses under “Input Image” indicate input image shape (height, width, BGR color channels), while those of “Stomata Coordinates with Labels” correspond to (number of detected stomata, [xmin, ymin, xmax, ymax, object confidenceness score, class index]). See Materials and Methods for further details. (B) Representative output of a processed image selected from the test dataset. Red and blue boxes represent detected coordinates of open and closed stomata, respectively. The red polygon indicates the detected area of the stomatal pore. Enlarged views of detected stomata are displayed on the right.

To train the machine learning models used in the pipeline, we created an annotated dataset of Arabidopsis stomata from leaf disc images captured with a brightfield microscope. To obtain various appearances of stomata and mesophyll cells and enhance the general performance of our pipeline, we exposed plants to various external stimuli (different light conditions and chemical treatments) to affect the degree of stomatal opening (see Materials and Methods for details). We collected 79 images, resulting in the annotation of 1,300 open stomata and 880 closed stomata as bounding box coordinates, as well as 1,300 stomatal pore areas as polygon coordinates. We then separated these annotated images into training, validation, and test datasets (Fig. 2A). Representative results of the annotated images are displayed in Fig. 2B.

**Figure 2.**
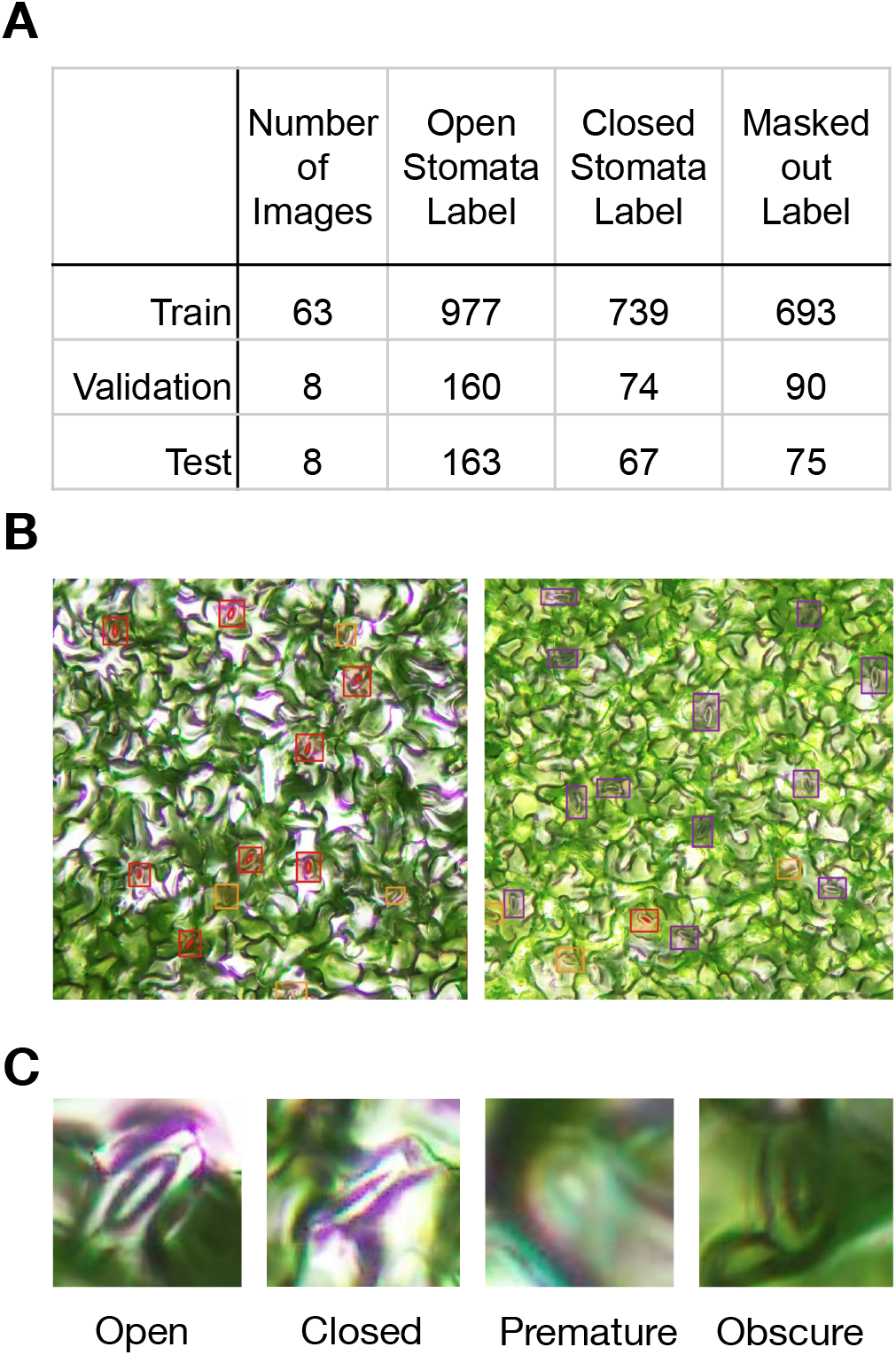
Dataset information. (A) Details of the annotated dataset of Arabidopsis leaf discs captured with a brightfield microscope. (B) Representative labeled images used in this study. Red and purple boxes represent labeled coordinates of open and closed stomata, respectively. The red polygon indicates the labeled area of the stomatal pore. Light blue and orange boxes represent premature stomata or stomata with obscure appearance, respectively. (C) Enlarged image of representative stomata from the four labeled categories.

We noticed a number of stomata with an ‘obscure’ appearance when they were out of focus or partially occluded by the surrounding leaf surface structure (Fig. 2C, obscure). We also observed the visuals of developmentally premature stomata suffering for the same reasons (Fig. 2C, premature). Such ‘noisy labels’ affected the quality of our manual annotation by mislabeling open or closed stomata, mislabeling non-stoma background features, or even incorrectly tracing stomatal pore perimeter. Since such low-quality features influence the performance of the machine learning model, these labeling issues must be addressed. Here, instead of repetitive manual data cleansing, we opted to omit any structure not clearly annotated as stomata and masked the corresponding areas as black rectangles resulting in labels of visibly clear stomata (Fig. 2C, open and closed). See Materials and Methods for details of the dataset generation.

For the stomatal detection model, we used YOLOX (You Only Look Once X) (Ge et al., 2021), a deep neural network aimed for object detection. We trained six YOLOX architectures (YOLOX-nano, -tiny, -s, -m, -l, and -x) with two input image size conditions (1280 × 1280 and 1920 × 1920). After training, we then compared the detection metrics against the test dataset. Among the trained models, YOLOX-x and YOLOX-s with the 1920 × 1920 input image size displayed the highest mAP_50_ (mean average precision) values of 0.930 and 0.875, respectively (Fig. 3A). Considering that mAP_50_ values for both models were higher than 0.85, we selected YOLOX-s with 1920 × 1920 input image size for the pipeline, which has a smaller network architecture, to prioritize pipeline processing speed. For the isolation of stomatal pores, we separately implemented two semantic segmentation model architectures (U-Net and DeepLabv3) (Chen et al., 2017; Ronneberger et al., 2015) each with two types of encoder backbones (MobileNetV3-Small and MobileNetV3-Large) (Howard et al., 2019). Out of the four combinations tested, the U-Net architecture with the MobileNetV3-Large backbone returned the highest IoU with a value of 0.745 (Fig. 3B). We thus selected this combination for the segmentation submodule of our pipeline. These detection and segmentation models along with post-processing steps resulted in a stomatal aperture quantification error compared to manual measurement (ground truth) of 0.2 ± 0.2 μm (see Supplementary Fig. 1 for measurement error on all test images) on average in the test dataset. Pipeline-processed results (prediction) and hand-annotated results (ground truth) were comparable in each image, regardless of the sample condition (Fig. 3C). Notably, the masking out of noisy labels had a limited effect upon such results (prediction with mask/prediction without mask). We conclude that we established an automated pipeline for the quantification of stomatal aperture from Arabidopsis leaf disc images.

**Figure 3.**
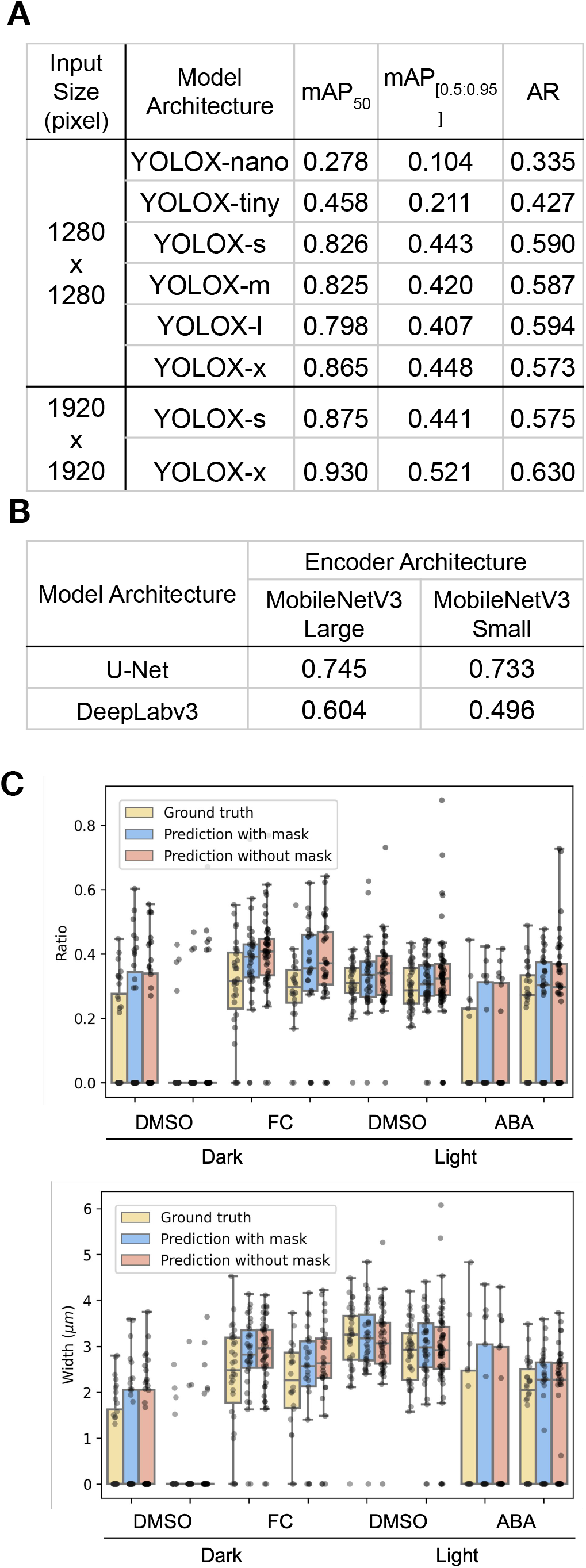
Pipeline performance. (A) Object detection metrics, for model architectures of trained stomata detection against the test dataset when masking noisy (premature and obscure stomata) labels. mAP50, mean average precision (mAP) with IoU threshold of 50%. mAP_50:95_, average of mAPs of IoU threshold from 0.5 to 0.95 with the step size of 0.05. AR, average recall. (B) Semantic segmentation metrics (intersection of union, IoU) for model architectures of the trained stomatal pore segmentation against the test dataset. (C) Graphs showing stomatal aperture (top, ratio; bottom, width) values measured either by hand-measured ground truth (yellow), prediction with mask (blue), or prediction without mask (pink) against eight images of the test dataset. Samples were either treated with 50 μM FC, 10 μM ABA, or DMSO (mock) and incubated under dark or Red-Blue light. See materials and methods for details.

### Portable Stomatal Imaging Device

While the above-mentioned models in the pipeline were trained and intended for use with images collected from isolated leaf discs with a benchtop microscope, we wished to expand the use of our pipeline to observe and quantify stomatal aperture from both detached and undetached intact leaves on-site. Therefore, we designed a portable device that can acquire the stomatal images by pinching the leaf blade with the help of image acquisition software (Fig. 4A, left). This device allows for the non-destructive (or less invasive) imaging of stomata. We reached an image sampling rate of about 0.5 μm/pixel with an optical resolution of 1 μm (see Materials and Methods for details). However, the model that was trained with microscopy images from leaf discs did not show sufficient detection and segmentation accuracy against images taken with the device, prompting us to fine-tune the trained model with an additional dataset acquired by this device until sufficient results were obtained (See Supplementary Fig. S3 and Materials and Methods for details). The use of this device with the fine-tuned model enables the precise acquisition of stomatal images and analysis, exemplified by the visualized result of undetached intact leaves (Fig. 4A, middle and right column).

**Figure 4.**
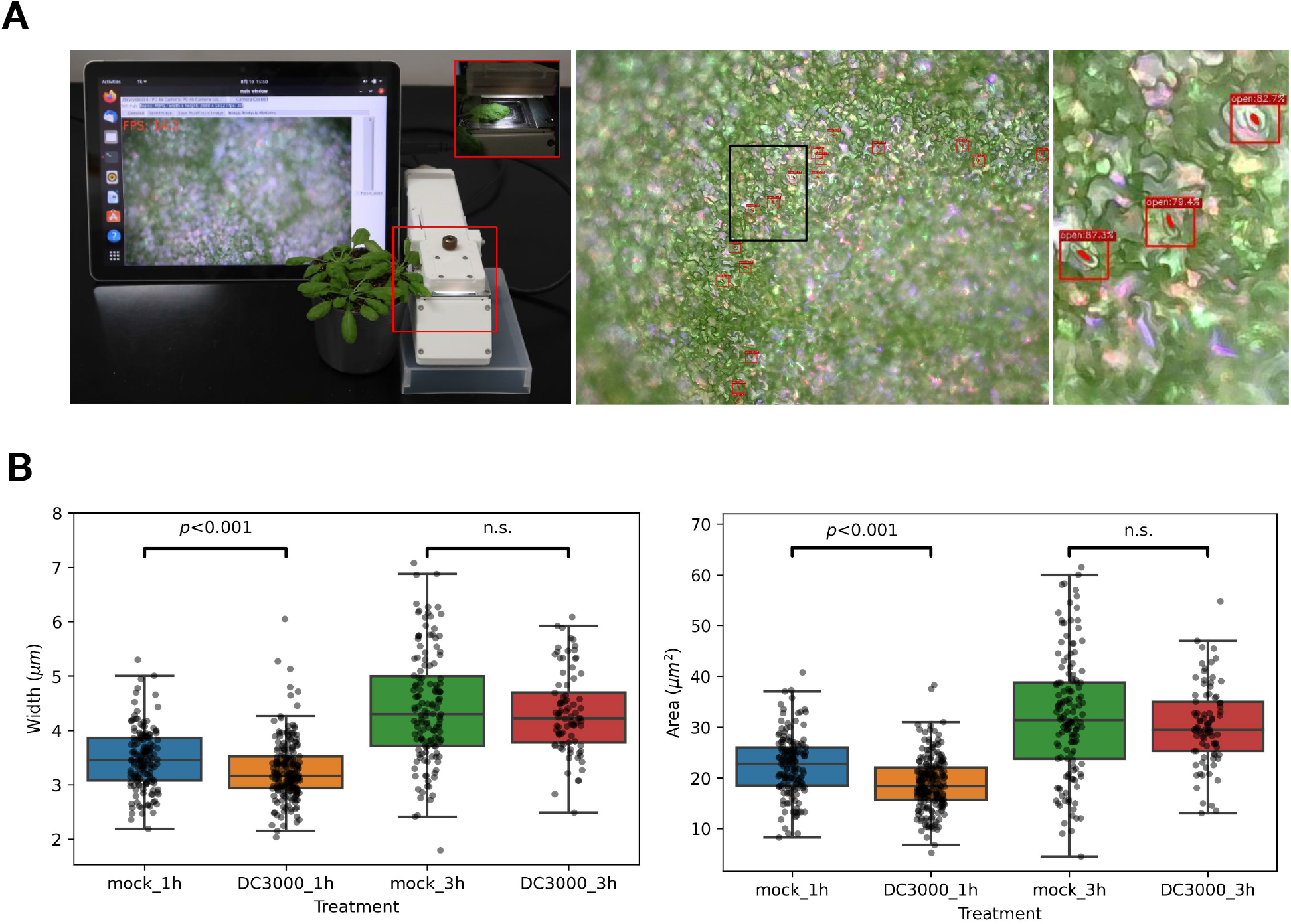
Portable imaging device. (A) Left, appearance of the portable device. Red insets show the mechanism pinching leaves without leaf detachment. Middle, representative result of an undetached intact leaf image acquired by the portable device, processed by the pipeline for stomatal aperture quantification. The area highlighted by the black rectangle is enlarged to the right. (B) Stomatal aperture (width and area) obtained by the pipeline. Detached leaves were immersed with a buffer alone or containing *Pst* DC3000, and were incubated for 1 or 3 h before the images were acquired, and stomatal apertures were measured. p value indicates the value of student t-test. n.s. indicates no significant difference between the two conditions. The same test images were used in the manual measurement of the stomatal aperture described in Supplementary Fig. S4.

Using the fine-tuned pipeline and the portable device, we investigated the response of Arabidopsis stomata against inoculation with *Pst* DC3000. According to the previously established method (Melotto et al., 2006), abaxial sides of detached leaves were placed onto stomata opening buffer alone or that containing bacteria, and stomatal images were collected with the stomata imaging device at 1 and 3 h after inoculation. Manual measurement of stomatal aperture confirmed previously reported stomatal responses to *Pst* DC3000; stomata close at 1h but open at 3 h (Supplementary Fig. S4) (Melotto et al., 2006). Prediction of stomatal aperture by our fine-tuned pipeline nicely recapitulated the stomatal responses to *Pst* DC3000 (Fig. 4B). Notably, the method introduced in the prior study required peeling of the epidermal layer from the detached leaves prior to microscopic observation, followed by manual measurement. Image acquisition without such sample preparation and automatic analysis of the pipeline obviously allows conduction of the experiment at higher throughput. We conclude that the device we present here now makes it possible to easily observe the stomata of leaves on-site.

## Discussion

In this study, we implemented an image analysis pipeline intended for quantifying stomatal aperture from Arabidopsis leaf disc images. We also developed a portable device specialized for easy image acquisition of leaves on-site. Utilizing such a device and/or the pipeline should enable researchers to accelerate routine analysis of stomatal aperture for physiological dissection of model dicot species.

Recent physiological research involving quantification of plant stomatal aperture greatly relies on isolated epidermal peels or leaf discs. Any interpretation of the derived results must therefore take into account unavoidable effects of phytohormones such as jasmonate and ABA on the experimental results, as their biosynthesis is triggered through sample preparation (e.g., physical damage and water loss). Ideally, utilizing undetached intact leaves in any situations should enable observation of the ‘natural’ physiological response of plants, repetitive analysis of the same plantlet at different time points, and analysis of the effect of long-distance (leaf to leaf) signaling, such as defense signaling (Mousavi et al., 2013). Nonetheless, observations of stomata in epidermal peels, excised leaf discs, or detached/undetached intact leaves have their specific merits and are complementary approaches to the same end; they should be the subject of careful selection as a function of the research inquiry at play. Here, we assembled an analysis pipeline that can quantify stomatal features from images of leaf disks captured with a brightfield microscope and then adapted the training model to accommodate the use of images from intact leaves taken by a portable device to address unchallenged issues up-to-date.

Training machine learning models, specifically supervised learning, generally require an annotated dataset that consists of images and their corresponding object labels. For example, general object (e.g., a bus, a person, or a horse) detection and segmentation datasets such as COCO (common objects in context) were created by defining a bounding box and outline around a given object (Lin et al., 2014). The bounding box coordinates for the location of each stoma and the stomatal pore outline corresponds to such labels in our stomatal dataset. While COCO was annotated by non-experts hired through a crowd labeling service, our images were annotated by multiple experienced plant physiologists who, unlike non-experts, were trained to pinpoint ambiguous stomata “hiding” in brightfield images. Notably, we further tried to enhance the quality of our labeling by applying focus stacking (see Material and Methods) and masking noisy labels in the dataset. Nonetheless, we noticed inconsistent annotations from one expert to another even when processing the same image, although all images and annotations were double-checked by an independent expert (Supplementary Figure. S2). Such differences were likely derived from variations in bounding box sizes, false negative labels, and personal decisions on what constituted a premature or an obscure stoma. Ideally, such inconsistent criteria should be unified during annotation. However, contrary to general object detection tasks (i.e., labeling regions with a dog or a cat), reaching a consensus for stomatal boundaries and label classification requires a substantial amount of time. Rather than striving for perfection, we limited the extent of annotation refinement and trained our machine learning models in advance to verify that the quality of our dataset was sufficient for deployment. We anticipate that coming to terms with the quality of each dataset labeling will be integral to deploying the analysis pipeline when the objects of interest are more difficult to annotate.

The machine learning model trained on images acquired from leaf discs with a brightfield microscope was not directly compatible with our portable device, thus necessitating further fine-tuning (Supplementary Fig. S3). We suspect that this lack of initial compatibility was due to intrinsic differences in the acquired images derived from hardware configuration (imaging sensors, light intensity, and/or light color). Further analysis is needed to identify the cause, which is a common issue in the machine learning domain that is often referred to as extrapolation. Machine learning models cannot properly process a type of images that differ from that used for model training. A general solution is to simply generate a larger training dataset in the context of both image quantity and types of images acquired in various environments (e.g., type of image acquisition apparatus) to enhance the general applicability of the trained model. We anticipate that ongoing collections of various stomatal images will clarify these issues in future studies. Also, our trained pipeline was designed for a fixed image acquisition condition (see Materials and Methods for details) and is not a generalized stomatal aperture processing module. Importantly, our pipeline is not intended to quantify stomata from species other than Arabidopsis. Nonetheless, our approach utilizes generalized neural network architecture and is easy to adapt to a specific experimental condition, including another plant species, provided an appropriate training dataset is generated.

Here, we introduced an intuitive example of the utility of our stomatal aperture measurement pipeline and portable device by quantifying the stomatal response of Arabidopsis leaves to various environmental stimuli. Observation of stomatal response against chemical treatment in leaf discs (Fig. 4C), and pathogen inoculation against detached leaves are no more than a representative use case. For example, screening for Arabidopsis mutants, compounds, and pathogens that respectively influence the stomatal regulation is a potential scope.

Further investigation using this pipeline/device is expected to accelerate research aimed at elucidating the molecular mechanisms underlying stomatal regulation in response to various external stimuli, including light regulation and plant–microbe interactions.

## Materials and Methods

### Dataset Generation

Images used for generating the training dataset for the machine learning model were acquired as described previously (Toh et al., 2018). Briefly, *Arabidopsis thaliana* plants (Col-0 accession) were grown on soil at 22°C under a 16-h-Red/Blue-light (50 and 10 μmol/m^2^/s, respectively)/8-h-dark photoperiod. Prior to the day of image acquisition, plants were moved to a dark room for at least 12 h to induce stomatal closure. Under dim light, leaf discs were excised from the center of fully expanded rosette leaves using a hole puncher (6-mm-diameter Biopsy Punch; Kai Medical). Samples were then immersed in a basal buffer (5 mM MES-Bis-tris propane pH 6.5, 50 mM KCl, and 0.1 mM CaCl_2_) either containing 20 μM abscisic acid (ABA), 10 μM fusicoccin (FC), or 50 μM DMSO, and then further incubated in the dark or white light (50 μmol/m^2^/s) for 3 h. In this study, four conditions were selected to obtain images containing stomata in various conditions; ABA + light, DMSO + light, FC + dark, and DMSO + dark. Images were acquired using an upright optical microscope (BX43; Olympus, Tokyo, Japan) with a 10× objective lens with a CCD camera (DP27; Olympus) at a resolution of 1920 × 2448 (height × width) pixels and a scale of 0.35 μm/pixel using cellSens software (Olympus). The Instant Extended Focal Imaging (Instant EFI) add-on of cellSens was used upon image acquisition to gain as full a focus of the acquired view as possible.

To annotate images used for training the stomatal aperture quantification model, we utilized a cloud labeling service, Labelbox (Labelbox, 2022. https://labelbox.com). Four bounding box label classes were defined to mark “open stomata,” “closed stomata,” “premature stomata,” and “obscure stomata”, and an independent polygon label was used to trace the outline of stomatal pores. After labeling, datasets were converted to COCO format and used as the basis for the stomatal detection dataset. Bounding box coordinates of obscure stomata and premature stomata in the images were black-filled and omitted from the annotation. Upon model training in the later stage, datasets were divided into training/validation/test subsets. To prepare images for the stomatal aperture pore segmentation model, the center point of the bounding box was first calculated. Then, with its coordinate as a center, a subset image of 64 × 64 pixels was cropped along with the corresponding stomatal pore mask image to create a dataset.

### Image Analysis Pipeline

Stomata detection models were trained using YOLOX (Ge et al., 2021) following the manual of the official repository (https://github.com/Megvii-BaseDetection/YOLOX) with slight modifications. The dataset used for training was generated as described in the previous section. Prior to model training, five augmented images per image were generated using the *Rotate* and *ColorJitter* functions from the Albumentations library (https://github.com/albumentations-team/albumentations). The training was performed at an image resolution of 1280 × 1280 pixels or 1920 × 1920 pixels with 300 epochs. The batch sizes were set to 4 and 2 for the YOLOX-l and YOLOX-x models, respectively, and to 8 for the others for 1280 × 1280 input size. For 1920 × 1920 input size, batch sizes were set to 6 and 1 for the YOLOX-s and YOLOX-x models, respectively. After training, the model weight that displayed the best mean average precision (mAP50) against the validation dataset was selected.

Stomatal pore segmentation models were trained using the pytorch segmentation library following the procedure of the official repository (https://github.com/qubvel/segmentation_models.pytorch). Images were augmented upon training with the *HorizontalFlip* and *ShiftScaleRotate* functions and then *CLAHE, RandomBrightnessContrast, RandomGamma, IAASharpen, Blur, MotionBlur*, and *HueSaturationValue* functions randomly from the Albumentation library. Training was carried out with 400 epochs with a 64 × 64 pixel input size. After training, the model weight that displayed the best intersection of union (IoU) against the validation dataset was selected. Stomatal aperture quantification from stomatal pore mask images was performed by obtaining the minor axis length of the segmented region using the *measure.regionprops* module of the scikit-image library. Finally, the stomatal detection model and the stomatal pore segmentation model were converted from pytorch to ONNX format upon deployment.

Additionally, fine-tuning (i.e. further training of models to adapt to a new dataset) was performed against the above-mentioned detection and segmentation models with a dataset whose images were taken by the portable imaging device with detached/undetached intact leaves (Supplementary Fig. S3A). In contrast to the dataset of microscopic leaf disc images, the majority of the stomata were opened, and completely closed stomata were absent. This may be due to the condition of the sample or observation condition but reasons are currently unknown. Labeled stomata in the dataset were all handled as open stomata, and pore segmentation was performed against all objects. Similar to the training of microscopic models, stomatal detection and pore segmentation models were trained using YOLOX and the pytorch segmentation library by fine-tuning the model weights generated for the microscopy. For the stomatal detection models, five augmented images per image were generated using the *Rotate, ColorJitter, ShiftScaleRotate, Blur* and *HueSaturationValue* functions from Albumentations library. The training was performed at an image resolution of 1920 × 1920 pixels with 300 epochs. The batch sizes were set to 6 and 1 for the YOLOX-s and YOLOX-x models, respectively. For the stomatal pore segmentation models, 28 × 28 pixels cropped images were augmented with the *HorizontalFlip* and *ShiftScaleRotate* functions and then *CLAHE*, *RandomBrightnessContrast*, *RandomGamma*, and *HueSaturationValue* functions randomly from the Albumentation library, and finally resized to 96 × 96 pixels. The training was conducted with 400 epochs with 96 × 96 input image size. Upon inference, 28 × 28 of stomata images were resized to 96 × 96, inferred with the segmentation model, and resized back to 28 × 28, and further gone through the post processing for pore measurement. To measure the model performance, 32 images of detached leaves were additionally acquired and used as a test dataset. First, only the stomatal detection were performed, and were handed to the manual annotator to evaluate which detection were not suitable for biological analysis as well as identifying non-stomata detection (i.e. counting false positives). Out of 585 stomata detected by the model, 32 was manually judged as false positive, accounting to a precision value of 0.945. Then, stomatal pore segmentation with the analysis pipeline and manual-measurement with ImageJ were individually performed without containing false positives. The two sets of values were compared, plotted as a scatterplot, and pearson correlation were calculated (Supplementary Fig. S3C). This set of images were also used to generate Fig.4B, however the stomatal apertures deriving from the false positives were included to display the raw output values.

### Program Development Environment

Machine learning model training as well as pipeline construction were performed using a cloud programming service (Google Colaboratory Pro+) using Python language. All code development was done on a laptop computer (Macbook Air M1) or desktop computer (Mac mini 2018).

### Portable Imaging Device Design and Imaging Software

The hardware was designed to acquire stomatal images by pinching the plant leaf. The base unit was created using a filament PolyMAX PLA White (Polymaker, China) with an L-DEVO M2030 3D printer (Fusion Technology, Japan). An LED light (OSTCWBTHC1S; Optosupply Limited, China) irradiated downward through semi-transparent silicon rubber (GELS1-50; MISUMI, JAPAN) onto the adaxial side of the leaf. The silicon rubber was inserted to function as a light diffusion and a pressure source for sample flattening. A camera module (Happy Quality, Japan; CY&K INTERNATIONAL, Japan) was mounted to the bottom side of the device facing upward to acquire an image of the abaxial leaf surface. The camera module consisted of a compound of micro-size lens, focus control motor, and a complementary metal oxide semiconductor sensor (IMX 415; Sony, Japan), which can acquire 2592 × 1944 (height × width; pixels) images with a resolution of about 0.5 μm/px at maximum. A 32 × 32 mm cover glass (Iwanami, Japan) was inserted between the camera and the leaf. The device was connected to a computer by a USB 3.1 (GEN2) cable for data transmission and power supply. The camera module is UVC (USB Video Class) compatible so the input image can be displayed on any computer without a specific driver.

### Bacterial Inoculation

*Arabidopsis thaliana* plants (Col-0 accession) were grown on soil in a chamber at 22°C, under a 10-h-light/14-h-dark photoperiod and 60% relative humidity. Light intensity was abt. 5000 lx measured by an illuminometer, which corresponds to photosynthetic photon flux density of abt. 80 μmol/m^2^/s. Five-week-old plants were used for bacterial inoculation. To ensure that most stomata were open, plants were kept in the light for at least 3 h prior to assay. *Pst* DC3000 was cultured at 22°C in King’s B (KB) medium with 50 μg/mL rifampicin. The bacterial cultures were pelleted by centrifugation, washed twice with water, and resuspended in stomata opening buffer (25 mM MES-KOH pH 6.15, 10 mM KCl) (Melotto et al., 2006). Detached leaves were immersed in bacterial suspension at OD_600_ of 0.2 in stomata opening buffer and incubated under light at an intensity of abt. 10000 lx (abt. 170 μmol/m^2^/s).

## Data Availability

The model weights and codes in ONNX format to execute the Arabidopsis stomata quantification pipeline as well as mask and unmasked test data are available through (https://github.com/phytometrics/arabidopsis_leaf_stomata_quantification). The software used to process image streams from the portable device is under development at (https://github.com/phytometrics/cvgui_linux).

## Conflict of Interest

Y.T. is an employee of Phytometrics and Happy Quality. M. Y.-K. and M.T. are employees of Phytometrics.

## Author Contributions

M.Takagi, R.H., A.M., and Y.T, wrote the manuscript and got approval from all the coauthors. T.K., A.M., and Y.T. conceptualized this research. Project administration was done by Y.T.. M.Takagi and Y.T. developed the image analysis pipeline. Image acquisition and/or dataset labeling was performed by M.Takagi., R.H., Y.T., A.M., Y.A., Y.H., M.M.-A., E.A., M. Tomiyama, and M.K.-Y..

## Acknowledgements

We thank Dr. Kei Hiruma and Dr. Motoyuki Ashikari for sharing laboratory space and experimental instruments, and Dr. Takeshi Higa for assistance in preparing the plants used in this research. We also thank NST co. ltd. (Shizuoka, JAPAN) for assistance in designing the portable imaging device. Yoko Tomita assisted us in acquiring images for the Arabidopsis leaf disc dataset. We thank all the members of the project, “Co-creation of plant adaptive traits via assembly of plant-microbe holobiont” for fruitful discussions. This work was supported by Grant-in-Aid for Transformative Research Areas (21H05151 and 21H05149 to A.M. and 21H05152 to Y.T.).

---

**Supplementary Fig. 1.**
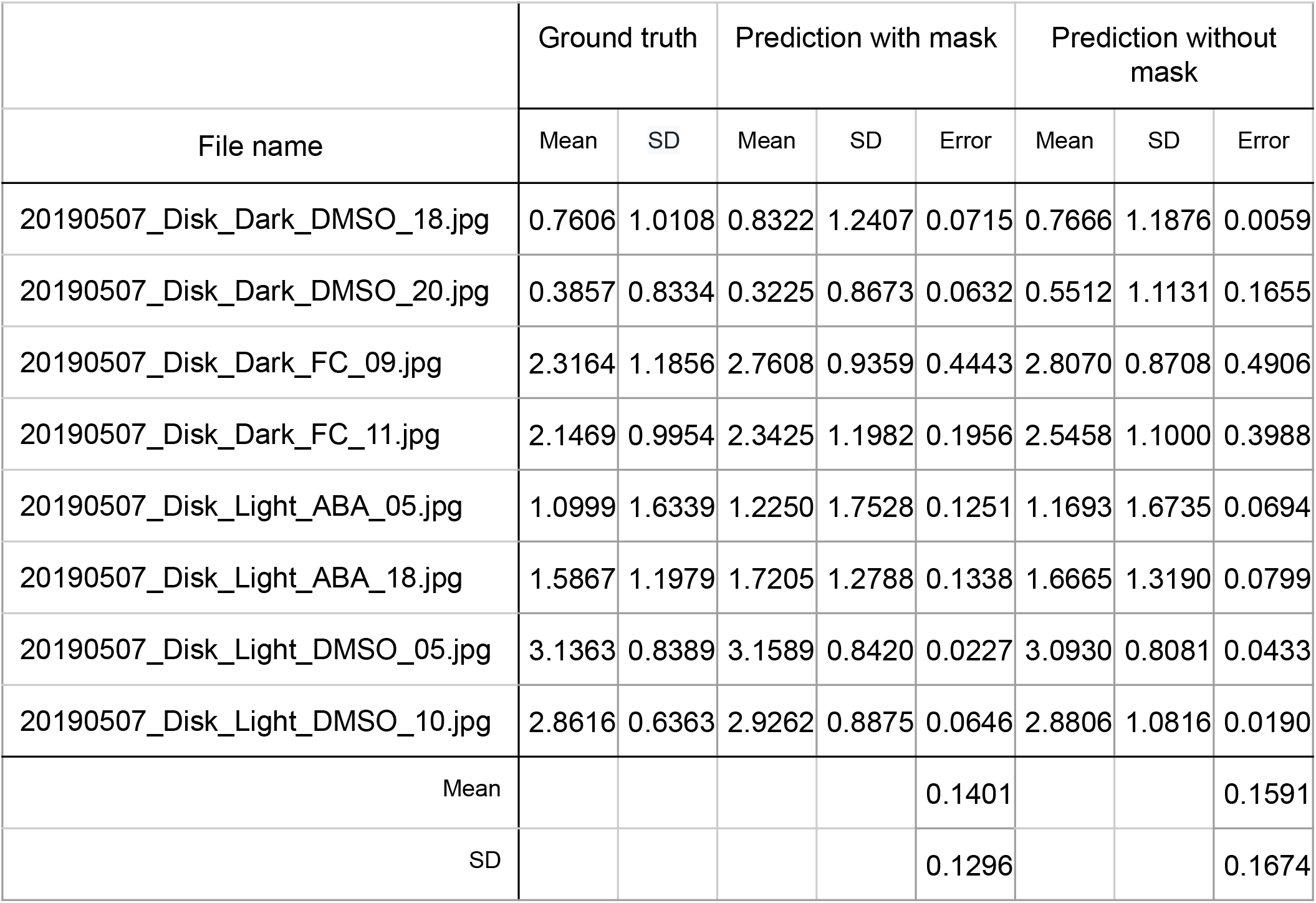
Stomatal aperture (width, μm) quantification difference between hand-measured ground truth and pipeline-processed results in test images. SD indicates standard deviation. Values of prediction with and without mask are displayed.

**Supplementary Fig. 2.**
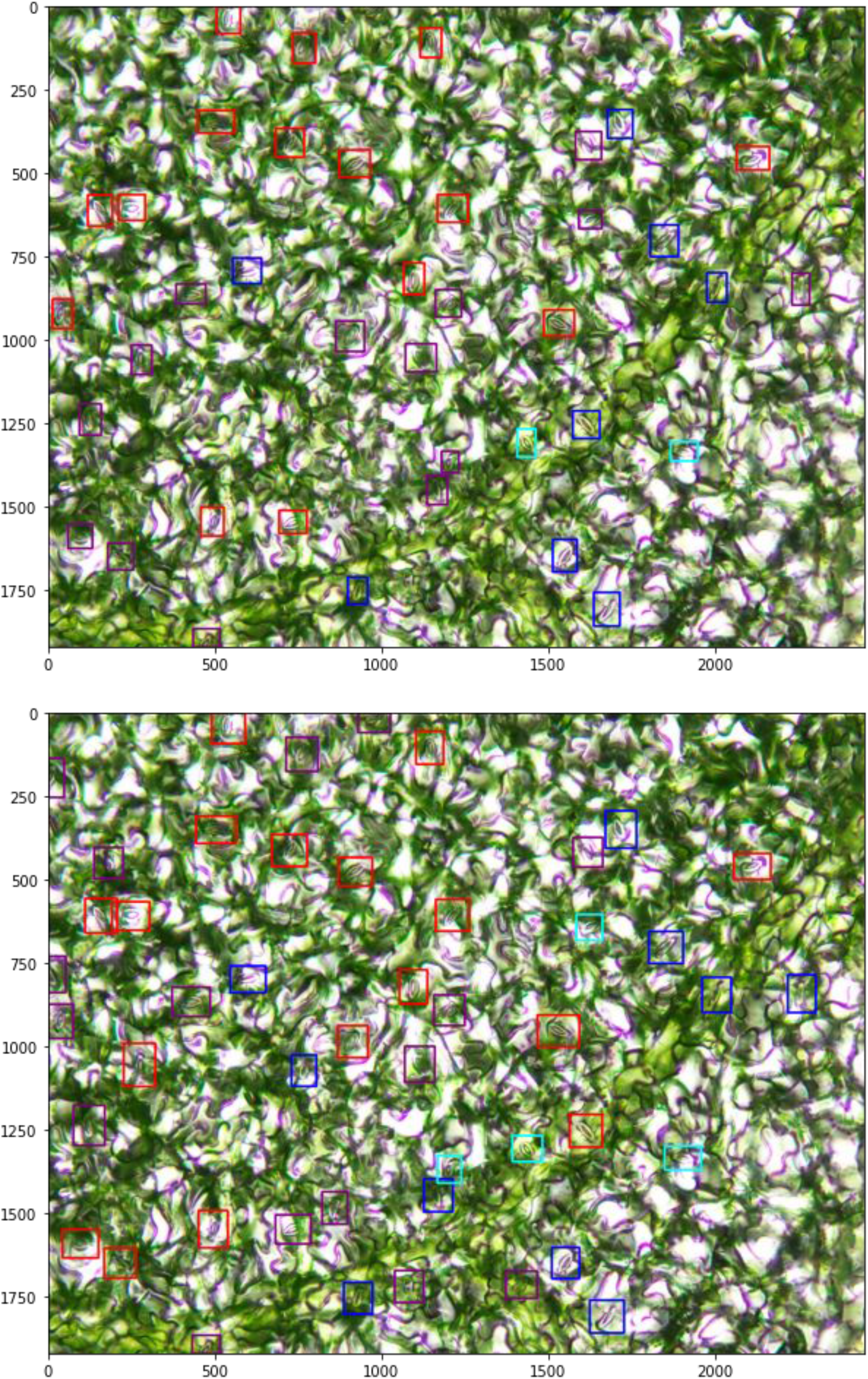
Representative result showing annotation inconsistencies. Top and bottom images were annotated by different labelers and reviewers.

**Supplementary Fig. 3.**
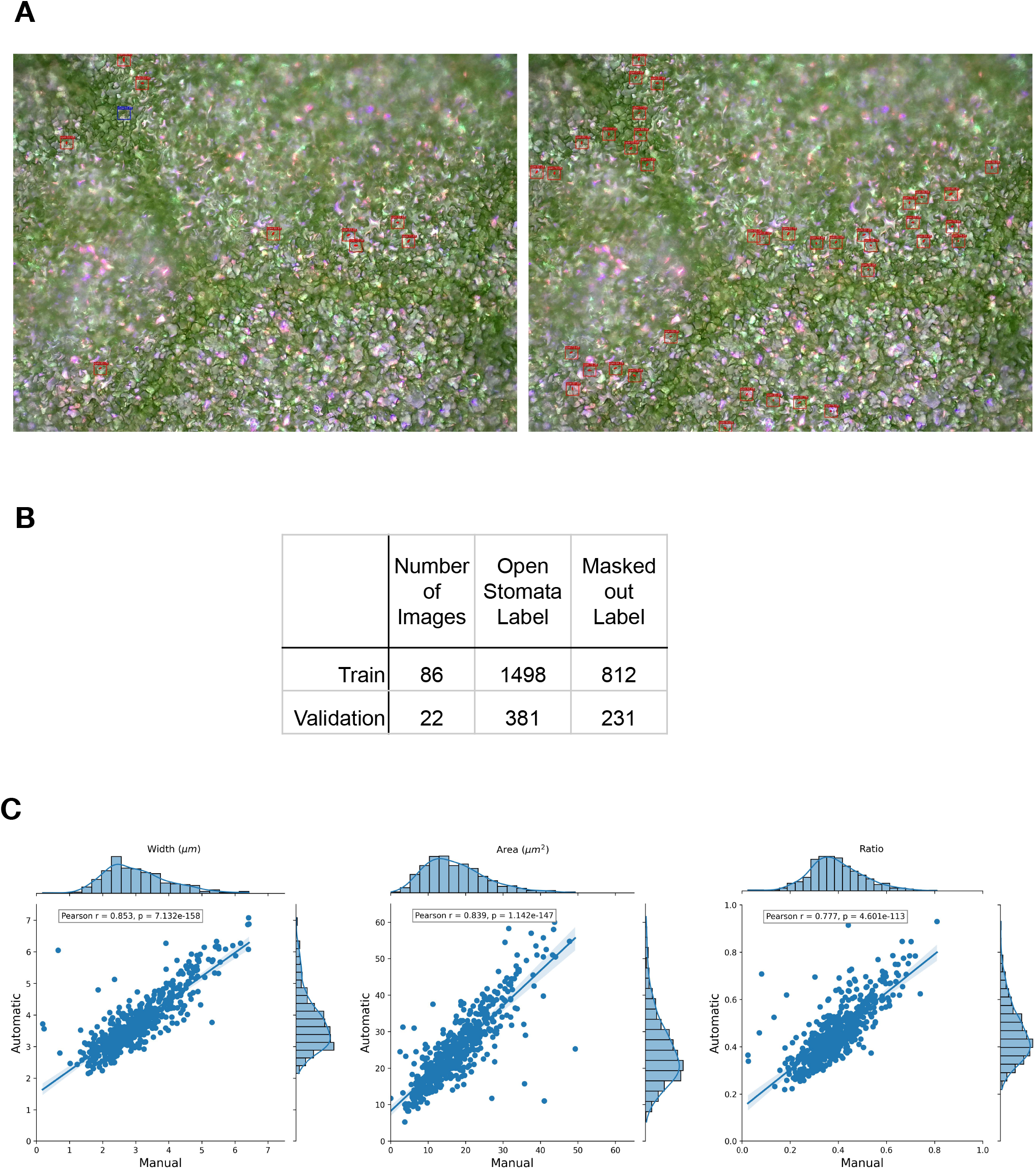
Adaptation of the pipeline suitable for the image acquired by the portable device. (A) Representative output of a processed image selected from the test dataset (same dataset used in Fig. 4B). Left, an image inferred by the models trained on images acquired by a brightfield microscope. Right, an image inferred by the models fine-tuned with images acquired by a portable device. (B) Details of the annotated dataset of Arabidopsis leaves captured with a portable device. (C) Scatter plots showing the correlation between manual measurement and automated-measurement stomatal aperture [Left, Width (μm); Middle, Area (μm^2^); Right, Ratio (Width/Length)] from the test dataset. The area values of manual measurement were calculated using the ellipse function in ImageJ. Pearson’s correlation coefficients (r) and p-values were indicated in the upper left of each panel.

**Supplementary Fig. 4.**
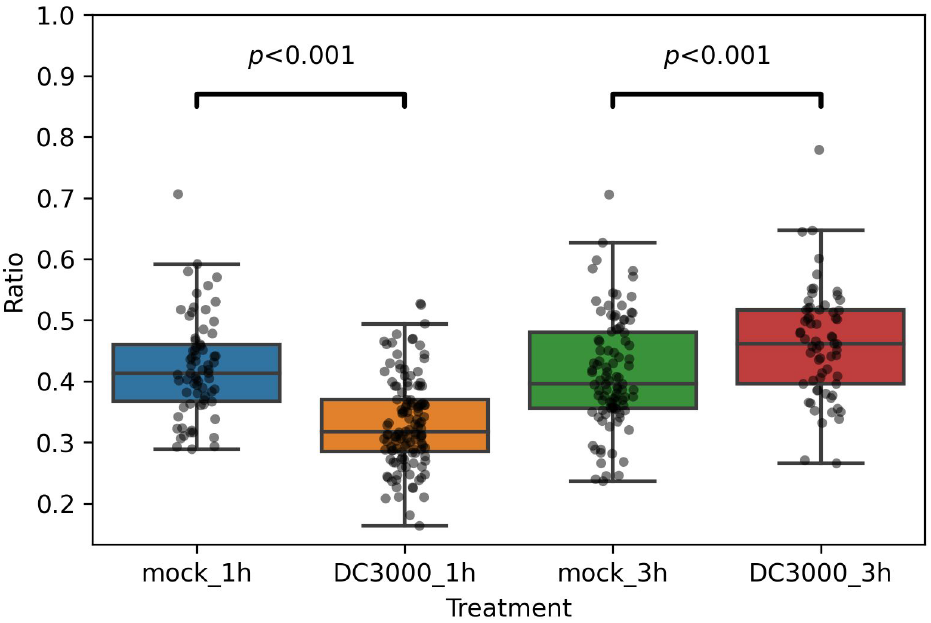
Manual measurements of Arabidopsis stomatal aperture (ratio) upon *Pst* DC3000 inoculation. In each experimental condition, at least 65 stomata were manually measured from two to three inoculated different leaves, and p-values were calculated by student’s t-test. This data shows the representative result of at least three independent experiments.

## Notes

https://github.com/phytometrics/arabidopsis_leaf_stomata_quantification

## References

Bourdais G, McLachlan DH, Rickett LM, et al. (2019) The use of quantitative imaging to investigate regulators of membrane trafficking in Arabidopsis stomatal closure. Traffic 20(2): 168–180. DOI: 10.1111/tra.12625.

Camoni L, Visconti S, Aducci P, et al. (2019) From plant physiology to pharmacology: fusicoccin leaves the leaves. Planta 249(1). Springer: 49–57. DOI: 10.1007/s00425-018-3051-2.

Chen L-C, Papandreou G, Schroff F, et al. (2017) Rethinking Atrous Convolution for Semantic Image Segmentation. arXiv [cs.CV]. Available at: http://arxiv.org/abs/1706.05587.

Clark D (2019) Stimulation and Observation of Leaf Stomata Using a Light Microscope. Microscopy today 27(4). Cambridge University Press: 18–23. DOI: 10.1017/S1551929519000622.

Eisele JF, Fäßler F, Bürgel PF, et al. (2016) A Rapid and Simple Method for Microscopy-Based Stomata Analyses. PloS one 11(10): e0164576. DOI: 10.1371/journal.pone.0164576.

Ge Z, Liu S, Wang F, et al. (2021) YOLOX: Exceeding YOLO Series in 2021. arXiv [cs.CV]. Available at: http://arxiv.org/abs/2107.08430.

Hack HRB (1974) The Selection of an Infiltration Technique for Estimating the Degree of Stomatal Opening in Leaves of Field Crops in the Sudan and a Discussion of the Mechanism which Controls the Entry of Test Liquids. Annals of botany 38(1). Oxford Academic: 93–114. DOI: 10.1093/oxfordjournals.aob.a084806.

Hashimoto Y, Ino T, Kramer PJ, et al. (1984) Dynamic analysis of water stress of sunflower leaves by means of a thermal image processing system. Plant physiology 76(1): 266–269. DOI: 10.1104/pp.76.1.266.

Higaki T, Kutsuna N and Hasezawa S (2014) CARTA-based semi-automatic detection of stomatal regions on an Arabidopsis cotyledon surface. Plant morphology 26(1): 9–12. DOI: 10.5685/plmorphol.26.9.

Howard A, Sandler M, Chu G, et al. (2019) Searching for MobileNetV3. arXiv [cs.CV]. Available at: http://arxiv.org/abs/1905.02244.

Hsu P-K, Dubeaux G, Takahashi Y, et al. (2021) Signaling mechanisms in abscisic acid-mediated stomatal closure. The Plant journal: for cell and molecular biology 105(2). Wiley: 307–321. DOI: 10.1111/tpj.15067.

Inoue S-I and Kinoshita T (2017) Blue Light Regulation of Stomatal Opening and the Plasma Membrane H+-ATPase. Plant physiology 174(2): 531–538. DOI: 10.1104/pp.17.00166.

Jayakody H, Liu S, Whitty M, et al. (2017) Microscope image based fully automated stomata detection and pore measurement method for grapevines. Plant methods 13: 94. DOI: 10.1186/s13007-017-0244-9.

Liang X, Xu X, Wang Z, et al. (2022) StomataScorer: a portable and high-throughput leaf stomata trait scorer combined with deep learning and an improved CV model. Plant biotechnology journal 20(3). Wiley: 577–591. DOI: 10.1111/pbi.13741.

Li K, Huang J, Song W, et al. (2019) Automatic segmentation and measurement methods of living stomata of plants based on the CV model. Plant methods 15. Springer: 67. DOI: 10.1186/s13007-019-0453-5.

Lin T-Y, Maire M, Belongie S, et al. (2014) Microsoft COCO: Common Objects in Context. In: Computer Vision – ECCV 2014, 2014, pp. 740–755. Springer International Publishing. DOI: 10.1007/978-3-319-10602-1_48.

Matthews JSA, Vialet-Chabrand S and Lawson T (2020) Role of blue and red light in stomatal dynamic behaviour. Journal of experimental botany 71(7): 2253–2269. DOI: 10.1093/jxb/erz563.

Melotto M, Underwood W, Koczan J, et al. (2006) Plant stomata function in innate immunity against bacterial invasion. Cell 126(5). Elsevier: 969–980. DOI: 10.1016/j.cell.2006.06.054.

Melotto M, Zhang L, Oblessuc PR, et al. (2017) Stomatal Defense a Decade Later. Plant physiology 174(2). academic.oup.com: 561–571. DOI: 10.1104/pp.16.01853.

Mousavi SAR, Chauvin A, Pascaud F, et al. (2013) GLUTAMATE RECEPTOR-LIKE genes mediate leaf-to-leaf wound signalling. Nature 500(7463): 422–426. DOI: 10.1038/nature12478.

Purwar P and Lee J (2019) In-situ Real-time Field Imaging and Monitoring of Leaf Stomata by High-resolution Portable Microscope. bioRxiv. DOI: 10.1101/677450.

Ronneberger O, Fischer P and Brox T (2015) U-Net: Convolutional Networks for Biomedical Image Segmentation. arXiv [cs.CV]. Available at: http://arxiv.org/abs/1505.04597.

Sai N, Bockman JP, Chen H, et al. (2022) SAI: Fast and automated quantification of stomatal parameters on microscope images. bioRxiv. DOI: 10.1101/2022.02.07.479482.

Schindelin J, Arganda-Carreras I, Frise E, et al. (2012) Fiji: an open-source platform for biological-image analysis. Nature methods 9(7): 676–682. DOI: 10.1038/nmeth.2019.

Schneider CA, Rasband WS and Eliceiri KW (2012) NIH Image to ImageJ: 25 years of image analysis. Nature methods 9(7): 671–675. DOI: 10.1038/nmeth.2089.

Shimazaki K-I, Doi M, Assmann SM, et al. (2007) Light regulation of stomatal movement. Annual review of plant biology 58. esalq.usp.br: 219–247. DOI: 10.1146/annurev.arplant.57.032905.105434.

Shimono M, Higaki T, Kaku H, et al. (2016) Quantitative Evaluation of Stomatal Cytoskeletal Patterns during the Activation of Immune Signaling in Arabidopsis thaliana. PloS one 11(7): e0159291. DOI: 10.1371/journal.pone.0159291.

Song W, Li J, Li K, et al. (2020) An Automatic Method for Stomatal Pore Detection and Measurement in Microscope Images of Plant Leaf Based on a Convolutional Neural Network Model. Forests, Trees and Livelihoods 11(9). Multidisciplinary Digital Publishing Institute: 954. DOI: 10.3390/f11090954.

Toda Y, Toh S, Bourdais G, et al. (2018) DeepStomata: Facial Recognition Technology for Automated Stomatal Aperture Measurement. bioRxiv. biorxiv.org. Available at: https://www.biorxiv.org/content/10.1101/365098v1.abstract.

